# Large DNA virus promoted the endosymbiotic evolution to make a photosynthetic eukaryote

**DOI:** 10.1101/809541

**Authors:** Mitsuhiro Matsuo, Atsushi Katahata, Makoto Tachikawa, Yohei Minakuchi, Hideki Noguchi, Atsushi Toyoda, Asao Fujiyama, Yutaka Suzuki, Takayuki Hata, Soichirou Satoh, Takuro Nakayama, Ryoma Kamikawa, Mami Nomura, Yuji Inagaki, Ken-ichiro Ishida, Junichi Obokata

**Author notes:** Correspondence: Junichi Obokata, Graduate School of Life and Environmental Science, Kyoto Prefectural University, Kyoto, 606-8522, Japan.

## Abstract

Chloroplasts in photosynthetic eukaryotes originated from a cyanobacterial endosymbiosis far more than 1 billion years ago^1-3^. Due to this ancientness, it remains unclear how this evolutionary process proceeded. To unveil this mystery, we analysed the whole genome sequence of a photosynthetic rhizarian amoeba^4^, *Paulinella micropora*^5,6^, which has a chloroplast-like organelle that originated from another cyanobacterial endosymbiosis^7-10^ about 0.1 billion years ago^11^. Here we show that the predacious amoeba that engulfed cyanobacteria evolved into a photosynthetic organism very quickly in the evolutionary time scale, probably aided by the drastic genome reorganization activated by large DNA virus. In the endosymbiotic evolution of eukaryotic cells, gene transfer from the endosymbiont genome to the host nucleus is essential for the evolving host cell to control the endosymbiont-derived organelle^12^. In *P. micropora*, we found that the gene transfer from the free-living and endosymbiotic bacteria to the amoeba nucleus was rapidly activated but both simultaneously ceased within the initiation period of the endosymbiotic evolution, suggesting that the genome reorganization drastically proceeded and completed. During this period, large DNA virus appeared to have infected the amoeba, followed by the rapid amplification and diversification of virus-related genes. These findings led us to re-examine the conventional endosymbiotic evolutionary scenario that exclusively deals with the host and the symbiont, and to extend it by incorporating a third critical player, large DNA virus, which activates the drastic gene transfer and genome reorganization between them. This *Paulinella* version of the evolutionary hypothesis deserves further testing of its generality in evolutionary systems and could shed light on the unknown roles of large DNA viruses^13^ in the evolution of terrestrial life.

## Main manuscript

Our laboratory culture of *P. micropora* MYN1^5,6^ (Fig. 1a) is not axenic and contains bacteria. From this culture, we prepared the chromatins of *P. micropora* by micromesh-aided cell isolation and chromatin immunoprecipitation using a canonical histone antibody. Shotgun sequencing of this chromatin DNA gave us a high-quality draft genome assembly of 967 Mb (Fig. 1, Extended Data Fig. 1, Supplementary Table 1). K-mer analysis estimated the genome size of *P. micropora* MYN1 to be 1.35 Gb (Extended Data Fig. 1); hence, our draft assembly covered 72% of the whole genome. The genome is largely composed of repeated sequences, with 19.2% unique sequences (Fig. 1c). Simple repeat sequences are extraordinarily rich, amounting to 19.6% (Fig. 1c, 1d). As much as simple repeats and transposons, 20.5% of the genome is occupied by unclassified repeat sequences that contain notable amounts of DNA virus-like fragments (Fig. 1c, Supplementary Table 2, 3).

**Fig. 1.**
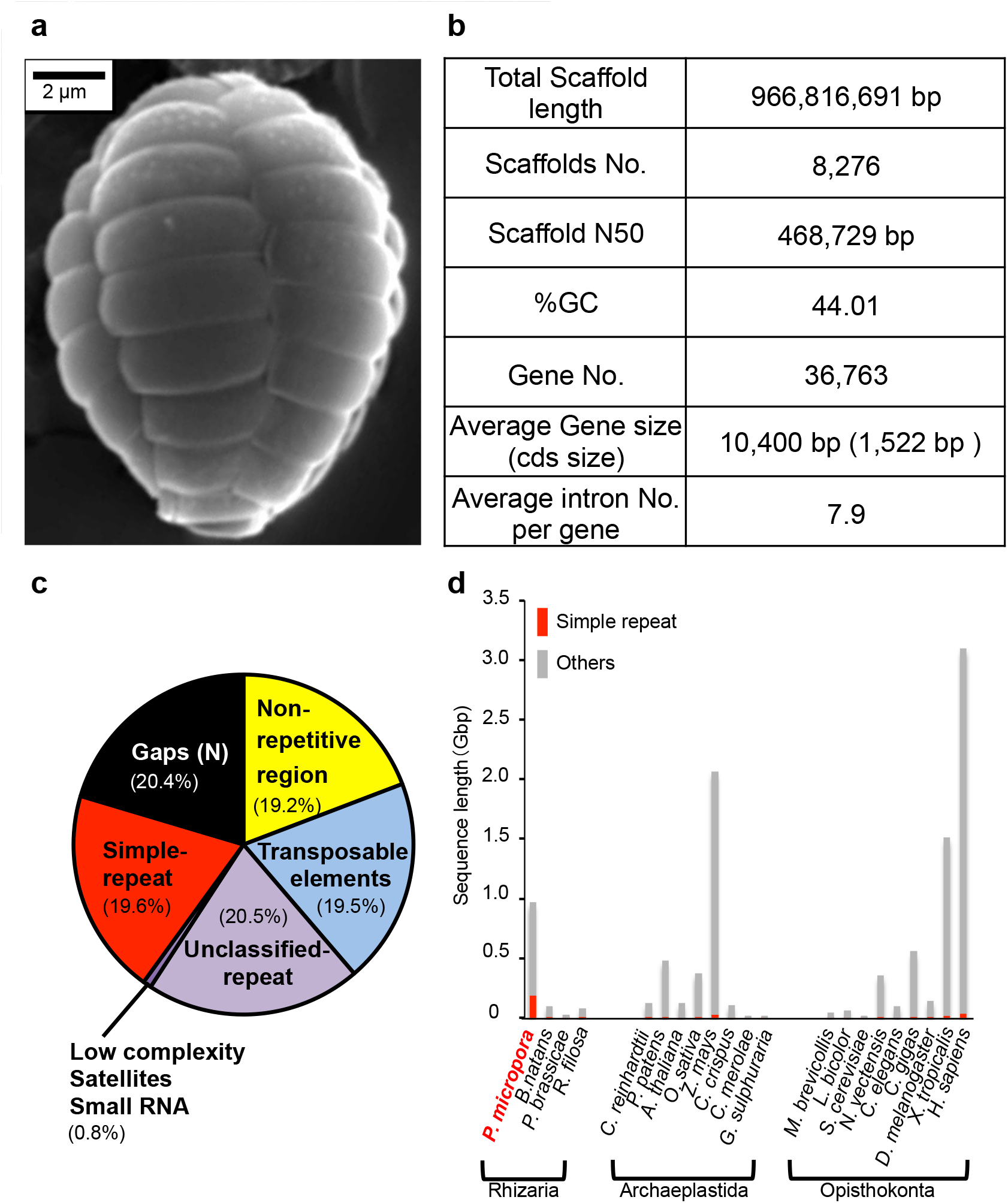
An overview of the *P. micropora* MYN1 draft genome. **a**, A SEM image of *P. micropora* MYN1. **b**, The statistics of the draft genome. **c**, The genome composition of *P. micropora* MYN1 analysed by RepeatMasker^27^. **d**, Simple repeats are extraordinarily rich in *P. micropora* MYN1 compared with other organisms.

A total of 36,763 protein gene models were predicted; on average, they were 10.4 kb long and contained eight introns, implying large and complex structures (Fig. 1b, Supplementary Table 4). Their gene ontology (GO) term analysis showed that DNA-related metabolism which associated with DNA virus is significantly over-represented compared with that of other rhizarian organisms (Extended Data Fig. 1e, Supplementary Table 5).

From the above gene set of *P. micropora*, we attempted to characterize the genes that have been pivotal for the endosymbiotic evolution. We extracted the genes derived from cyanobacteria as well as those derived from the rest of the bacteria; we refer to the former as endosymbiotic gene transfer (EGT) candidates and the latter as horizontal gene transfer (HGT) candidates in this study. We obtained 177 EGT and 248 HGT candidates (Fig. 2, Supplementary Table 6). Half of the EGT candidates are genes for high light inducible proteins (HLIPs)^12^, which are involved in the protection against excess light energy. Phylogenetic analysis of these HLIPs showed that they are polyphyletic, suggesting that HLIPs should have been acquired by multiple independent gene transfers from cyanobacteria (Extended Data Fig. 2). Thus, the gain of a light protection system should have been crucial for the predacious amoeba to evolve into a photosynthetic organism.

**Fig. 2.**
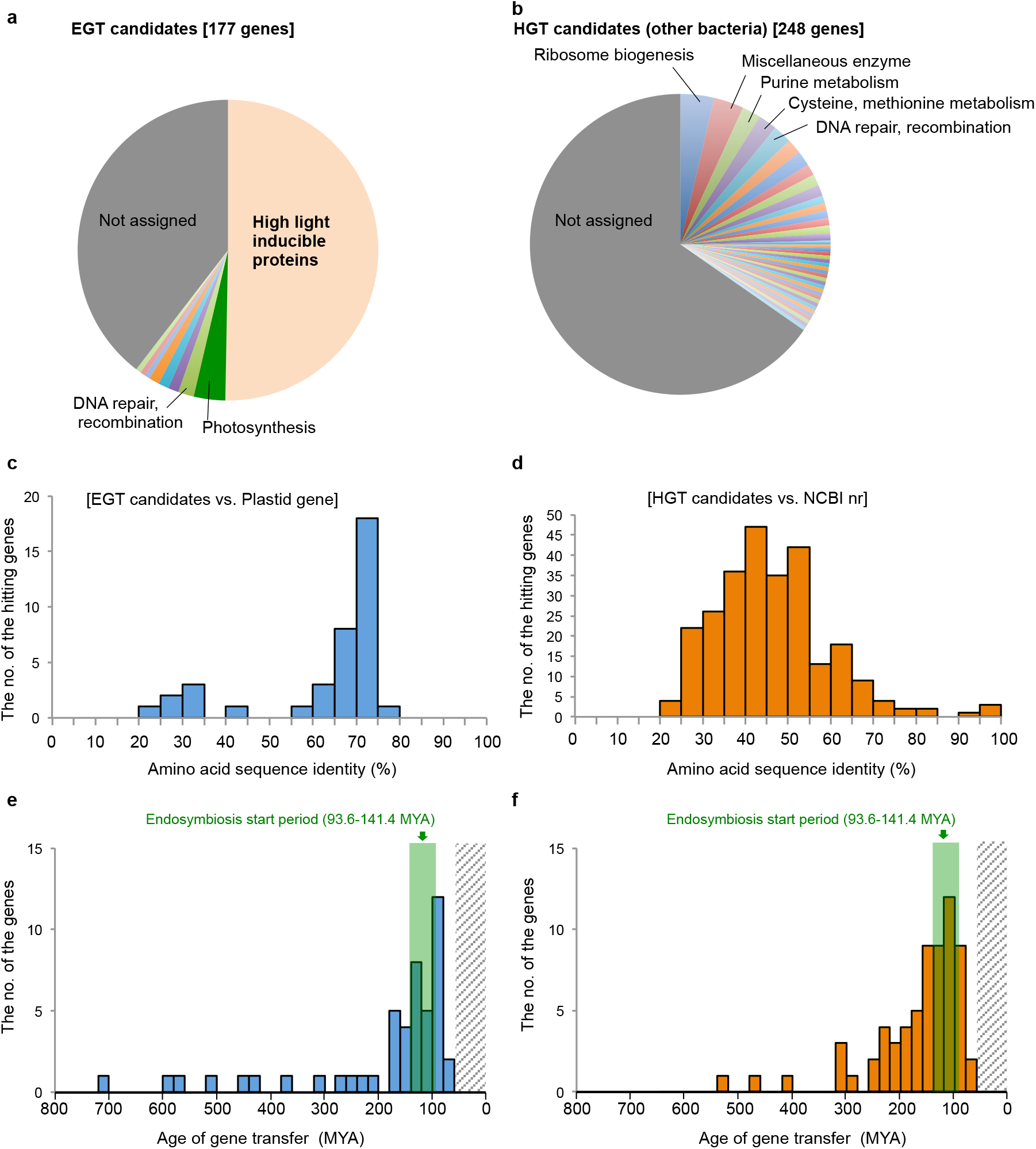
*P. micropora* nuclear genes acquired by EGT/HGT. **a, b**, A functional classification of the *P. micropora* nuclear genes derived from cyanobacteria (EGT candidates) (**a**), and those from other bacteria (HGT candidates) (**b**). **c, d**, The amino acid sequence identity of EGT candidates against *P. micropora* MYN1 plastid genes (**c**) and that of HGT candidates against bacterial genes of the NCBI nr database (**d**). **e, f**, An estimation of the gene transfer age for EGT candidates (**e**) and HGT candidates (**f**). The endosymbiosis initiation period is green-highlighted. The ages of gene transfer in (**e**) and (**f**) were calculated based on the divergent time points (45.7–64.7 MYA) of two *Paulinella* species; thus, a gene transfer age younger than 60 MYA (striped phase) could not be estimated.

HGT candidates contain genes of diverse functions, including ribosome biogenesis, DNA synthesis and amino acid metabolism. These genes appear to be involved in (1) endosymbiont biogenesis and (2) changes of the cellular nutrient state from heterotrophy to photo-autotrophy. To further examine the genes essential to the evolution of a photosynthetic organism, we compared orthologs among *P. micropora*, primary photosynthetic eukaryotes and predaceous eukaryotes (Extended Data Fig. 3a, 3b); 12 orthologous groups are conserved in the former two but not in the latter, including the genes for light acclimation, organelle gene expression and changes of the cellular nutrient state. Some of them were obtained horizontally from eukaryotes (Extended Data Figs. 3b–d). Therefore, *P. micropora* utilized the genes of diverse origins for endosymbiotic evolution.

**Fig. 3.**
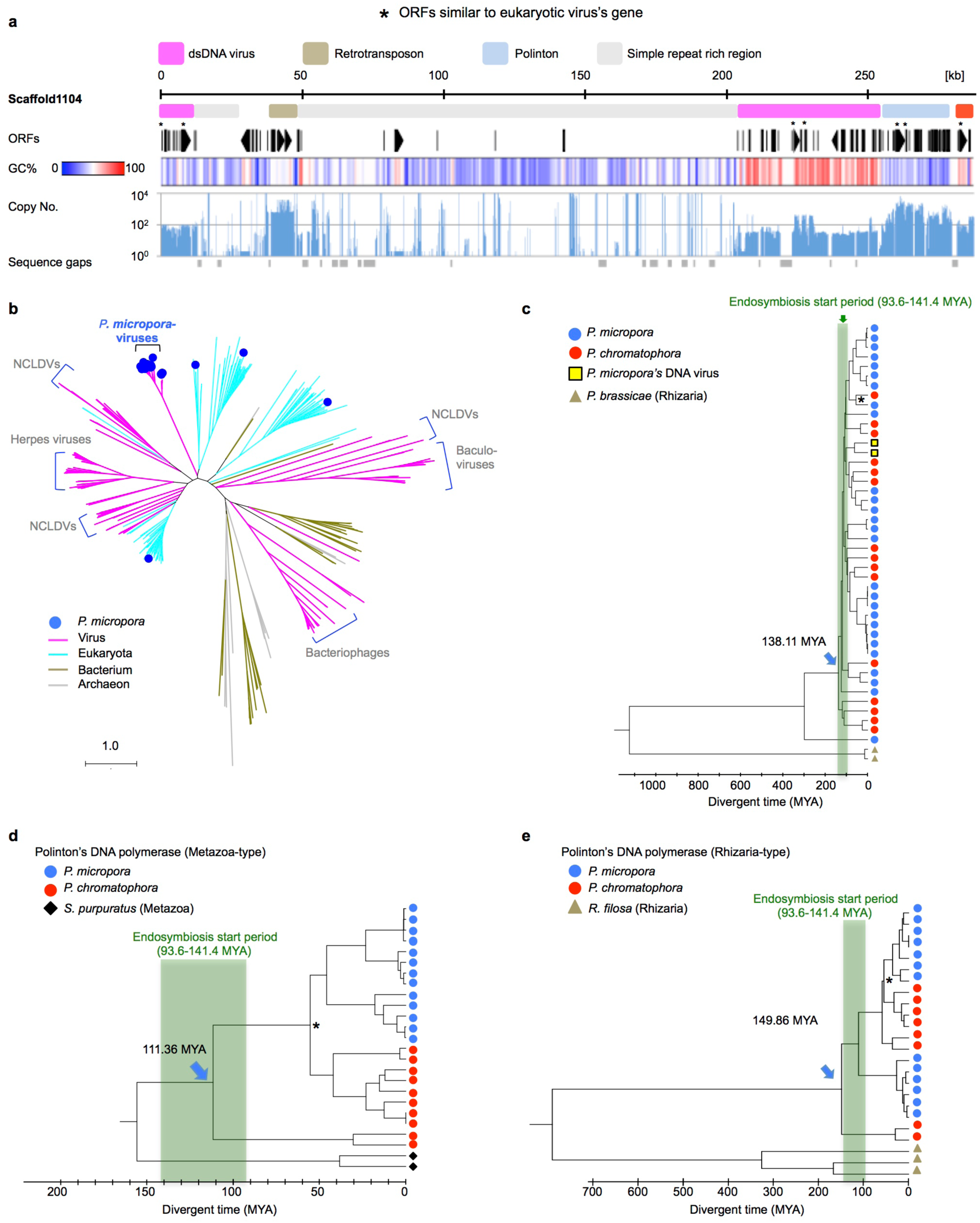
Putative DNA virus and mobile elements in *P. micropora* MYN1. **a**, A schematic view of DNA virus-like fragments and mobile elements in the *P. micropora* draft genome (Scaffold 1104). The genomic regions were coloured according to the sequence characteristics; putative dsDNA virus (pink), Polinton (light blue), retrotransposon (brass yellow) and simple repeat-rich region (grey). The copy number of the interspersed repeat elements was analysed by BLASTN against the simple-repeat-masked *P. micropora* draft genome. **b**, ML phylogenetic tree of DNA polymerases of viruses, eukaryotes and prokaryotes. **c**, Divergent time analysis of the virus-type GPCR in *Paulinella*’s lineage. **d, e**, Divergent time analysis of DNA polymerase genes of metazoa-type (**d**) and rhizarian-type (**e**) Polintons. Asterisks: the branch point of *P*. *micropora* and *P. chromatophora* set at 45.7–64.7 MYA. Green bands: initiation periods of endosymbiosis with cyanobacteria (93.6–141.4 MYA). *P. micropora*; *Paulinella micropora* MYN1, *P. chromatophora*; *Paulinella chromatophora* CCAC0185, *P. brassicae*; *Plasmodiophora brassicae*, *R. filosa*; *Reticulomyxa filosa*, *S. purpurgus*; *Strongylocentrotus purpuratus*.

The biggest challenge of this study is to elucidate the temporal sequence of the events that occurred at the birth of photosynthetic eukaryotes. To solve this puzzle, we first estimated how and when EGT occurred, based upon the sequence similarity between the EGT candidates and organelle-encoded genes. The results were surprising. We could not find any case with more than 80% amino acid sequence identity conserved between them (Fig. 2c), suggesting that plastidial EGT did not occur in a recent time period (Fig. 2e). We further searched for nuclear-localized plastid DNAs and nuclear-localized mitochondria DNAs^15,16^ in the genome and found the latter but not the former. Therefore, it is likely that plastidial EGT rapidly activated and then ceased early in the endosymbiotic evolution in *P. micropora*, while this cool down was not found for mitochondrial EGT (Extended Data Fig. 4).

**Fig. 4.**
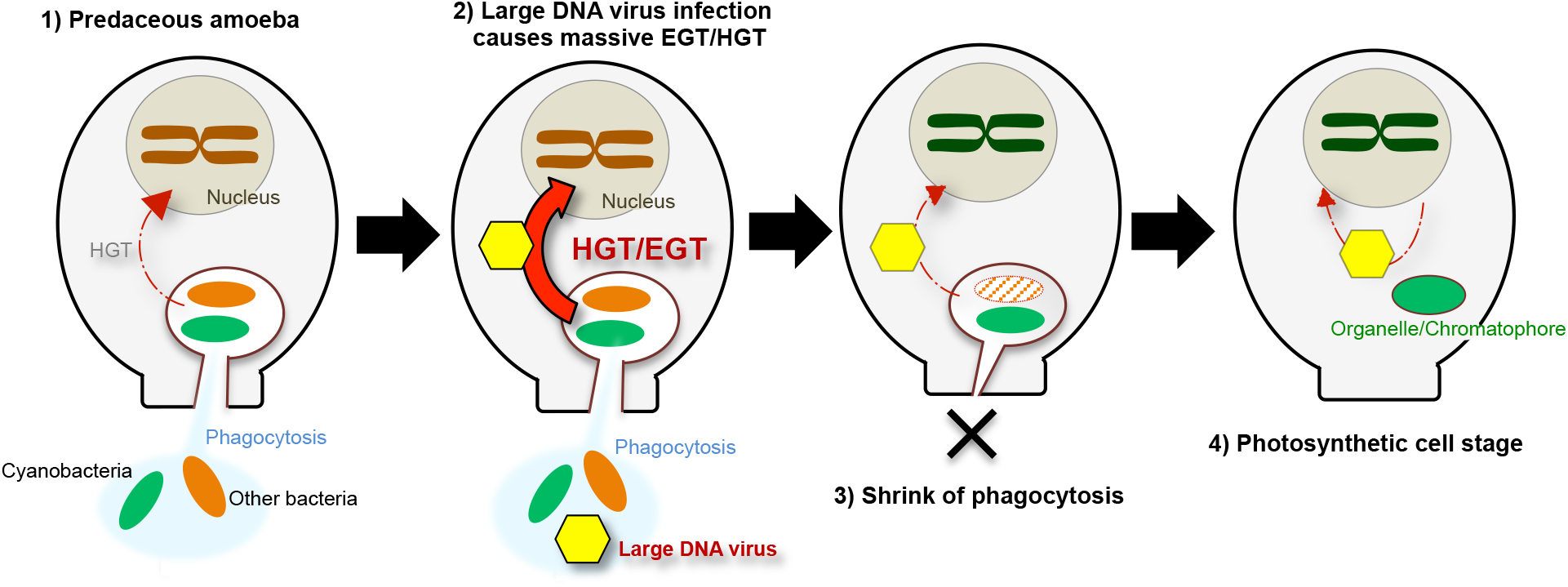
Initial process of the endosymbiotic evolution of photosynthetic *Paulinella* species modeled from the *P. micropora* genomic data. **1)** Predacious ancestors digested prey bacteria via phagocytosis with continuing low levels of HGT. **2)** The infection of large DNA virus triggered the massive HGT/EGT to promote the rapid endosymbiotic evolution. **3)** Acquiring photosynthetic competency shrunk the phagocytic activity to shut down the source of HGT/EGT. **4)** The photo-autotrophic *Paulinella* sp. contains one photosynthetic organelle per cell, hence, release of the organelle DNA hardly occurred without losing photosynthetic activity. In this final stage, virus-mediated gene transfer continued at trace level.

We confirmed this hypothesis from the different angle. Phylogenetic tree analysis of the EGT candidates showed that most of them already lost their counterparts in the plastid genome, except for four genes; hence, we reckon that these four genes were transferred from the plastid to the nucleus relatively recently. The divergences of the four genes between their nuclear and plastid counterparts were estimated to have occurred 319.8 to 98.6 million years ago (MYA) (Extended Data Fig. 5). Considering that the photosynthetic *Paulinella* species have diverged from the heterotrophic species 141.4 to 93.6 MYA^11^, even the latest EGT at 98.6 MYA had occurred within the initiation period of the endosymbiotic evolution. Taken together, the results of this study strongly suggest that EGT rapidly activated and ceased within the initial period of the endosymbiotic evolution (Fig. 2e), and a similar time course was also found for HGT (Fig. 2d, 2f).

What does this rapid and simultaneous cool-down of EGT and HGT (Figs. 2c–2f) mean? The most simple and likely explanation is that the predaceous *Paulinella* shrank and lost phagocytic activity at this time to become a photosynthetic organism, accompanied by the shut-down of phagocytosis-aided EGT/HGT. In reality, HGT from prey cyanobacteria occurred in the predaceous *Paulinella* species^17^. If our assumption is correct, the predaceous *Paulinella* should have changed its cellular, genomic and metabolic systems very quickly in terms of the evolutionary time scale. How was this drastic change possible? To examine this, we re-focused this study on DNA virus-like fragments frequently found in the *P. micropora* genome.

Fig. 3a shows a genomic scaffold containing putative virus fragments that are characterized by having from a dozen to a hundred copies, high GC content, ORFs similar to eukaryotic virus genes, and many intron-less genes of heterogeneous origins with unknown functions (Extended Data Fig. 6, Supplementary Table 3). In addition, they are often intermingled with simple repeats and mobile genetic elements, i.e., Maverick/Polinton-type giant transposons^18,19^ and retrotransposons. Most notably, the maximum fragment size reaches 300 kb (Extended Data Fig. 6). These structural features of the DNA virus-like fragments resemble those of nucleocytoplasmic large DNA viruses (NCLDV)^20^ whose genome size ranges from 100 kbp to 2.5 Mbp and who have many genes of heterogenous origins with unknown functions. However, a phylogenetic analysis based on DNA polymerases shows that those genes, encoded by the putative viral fragments, form a monophyletic clade distant from the genes of eukaryotes, prokaryotes and known NCLDVs (Fig. 3b). Therefore, we assume that they are from a novel large DNA virus but share several properties with known NCLDVs.

Our next question was when the putative virus infected the *Paulinella* lineage. Although ancient infection hallmarks were already smeared, we found a suggestive case in a *Paulinella*-specific gene family (Fig. 3c, Extended Data Fig. 7). The G-protein coupled receptor (GPCR) genes rapidly expanded and diversified within a short evolutionary period around the endosymbiosis initiation point. Noteworthily, two genes of this family were found only in the putative viral fragment regions (yellow squares in Fig. 3c and Extended Data Fig. 7). This suggests that these two genes diverged from the rest of the family around the endosymbiosis initiation point and have been inherited from the virus genome. This indicates that the putative virus has infected the *Paulinella* lineage around the endosymbiosis initiation point or earlier.

To further prove this, we investigated the Maverick/Polinton-type transposons derived from a virophage^21^, which parasitizes giant viruses (extremely large NCLDVs) with its propagation depending on the host virus^21,22^. Virophages are also integrated into the nuclear genome and could function as an anti-DNA virus system to protect eukaryotic cells from the DNA virus^23,24^. Therefore, we hypothesized that the emergence of Mavericks/Polintons and their amplifications have occurred concomitant with the DNA virus infection. In the *P. micropora* genome, two distinct Mavericks/Polintons, metazoan- and rhizaria-type, were detected in abundance (Extended Data Fig. 8, Supplementary Table 7). The divergent time analysis showed that both started amplification in the endosymbiosis initiation period (Fig. 3d, 3e). These results of the Maverick/Polinton-type transposons support the hypothesis that the large DNA virus infected the *Paulinella* lineage around the endosymbiosis initiation period.

Recent studies of NCLDV and giant DNA virus have drastically changed our conventional view of viruses^13^, especially their huge potential to incorporate diverse genetic materials of heterogenous origins^25,26^ and to mediate their shuffling. Considering these properties of large DNA virus and the results of this study, we could reconstruct the initial evolutionary process of the photosynthetic *Paulinella* species as shown in Fig. 4. In this hypothetical model, large DNA virus contributed to the endosymbiotic evolution as a critical player, in addition to the original players of the host and symbiont. We should note that, in general, the detection of ancient infection hallmarks of large DNA virus seems difficult because (1) it could be easily lost due to its harmful and undesirable effects on host proliferation, (2) of poor information of the virus sequences and (3) of repeated sequences that are apt to be omitted in the assembly process of the genomic sequencing projects. This *Paulinella* version of the endosymbiotic evolution hypothesis deserves further examination to test its generality in many evolutionary systems.

## Methods

### Data availability

The sequences of the *P. micropora* draft genome, plastid (chromatophore) genome, mitochondria genome and the raw reads data set were deposited to the DDBJ (Accession No. are shown in Supplemental Table 8) and DDBJ reads archives (DRA Accession No. DRA003059, DRA003106, DRA008524).

### *P. micropora* culture and cell isolation

The *P. micropora* MYN1 strain (NIES Collection, Tsukuba, JAPAN, NIES-4060) was cultured according to Nomura et al. (2014)^5^ and harvested by low-speed centrifugation (500 × g, 2 min) at 4 °C. The harvested cells were resuspended in the culture medium and filtrated through a 20 μm mesh nylon filter (HD-20, Nippon Rikagaku Kikai Co., Ltd., Tokyo, Japan) to remove dead cell aggregates with high bacterial contamination. Recovered healthy cells were repeatedly washed with culture medium and subjected to RNA extraction. For extraction of chromatin DNA, the cells were subsequently washed three times with 10 mM Tris-HCL (pH 8.0), six times with 10 mM Tris-HCL (pH 8.0) plus 10 mM EDTA, and recovered by a 5 μm mesh nylon filter (PP-5n, Kyoshin Rikoh Inc., Tokyo, Japan) to give clean cells largely free of bacterial contamination.

### Chromatin and genomic DNA extraction

Genomic DNA used for paired-end (300 b, 500 b) and mate-pair (3 kb, 5 kb) libraries for HiSeq sequencing were purified by chromatin immunoprecipitation (ChIP) as follows. Ten milligrams of *P. micropora* cells were homogenized in 500 μl homogenizing buffer (20 mM Tris-HCl (pH 7.6), 10 mM NaCl, 10 mM KCl, 2.5 mM EDTA, 250 mM sucrose, 0.1 mM spermine, 0.5 mM spermidine and 1 mM DTT) using a 30μm clearance glass homogenizer (RD440911, Teraoka Co., Ltd., Osaka, Japan). After centrifugation (1000 × g, 10 min, 4 °C), the pellets were resuspended in 300 μl ChIP buffer (50 mM Tris-HCl (pH 8.0), 500 mM NaCl, 10 mM EDTA, 0.1% SDS, 0.5% Na-deoxycholate, 1% Triton X-100, 1 mM DTT and 10% glycerol) with 20 μl Dynabeads protein G (Thermo Fisher Scientific, MA, U.S.A.) charged with 1 μg anti-histone H3 antibody (Ab1791) (Abcam plc, Cambridge, UK) and incubated at 4 °C for 20 min. Dynabeads were then washed twice with ChIP buffer, twice with glycerol-free ChIP buffer and finally suspended in DNA extraction buffer (10 mM Tris-HCl (pH 8.0), 1 mM EDTA and 1% SDS). After RNase A (10 μg/ml) and proteinase K (200 μg/ml) treatment, DNAs were purified by using Plant DNeasy Mini Kit (Qiagen, Hilden, Germany). For the construction of the long mate-pair libraries (12 kb, 15 kb, 18 kb and 20 kb), the total *P. micropora* genome was extracted without ChIP purification.

### Genome sequencing and assembly

Sequencing libraries were prepared using a TruSeq DNA PCR-Free Library Preparation Kit and a Nextera Mate Pair Library Prep Kit (Illumina, San Diego, CA). Two paired-end libraries with 300 and 500bp inserts and six mate pair libraries (3kb, 5kb, 12kb, 15kb, 18kb, and 20kb) were constructed and sequenced on the Illumina HiSeq 2500 sequencers with 151 cycles per run. The nuclear draft genome was assembled by SOAPdenovo v2.04-r240^28^ with a k-mer size of 121 after removing the sequence reads of the plastid (chromatophore) and mitochondria genomes, and those of two contaminating bacteria genomes. After the genome assembly, we checked for the contamination of the organelle genome and the bacteria genomes again, and we removed the contaminants from the draft genome. K-mer frequency analysis was performed by Jellyfish^29^. Genome scaffolds longer than 1 kb were analysed in this study.

### RNA-seq and Iso-Seq analysis

RNAs were extracted from *P. micropora* cells at 0, 4, 8, 12, 16 and 20 hr of 14L/10D photoperiod by Trizol® reagent (Thermo Fisher Scientific), and further purified using Plant RNeasy Mini Kit (Qiagen) with RNase-free DNase I treatment (Qiagen). Samples of the above time points were equally mixed and subjected to RNA-seq analysis using Agilent Strand Specific RNA Library Preparation Kit (Agilent Technologies, CA, U.S.A) and Illumina HiSeq 2500 (Illumina). The paired end reads of RNA-seq were *de novo* assembled by Trinity^30^ with the default setting or by CLC Genomics Workbench 7.0.3 (Qiagen) using a K-mer value of 54. The RNA-seq reads were mapped on the genome with Tophat^31^ and assembled with Cufflinks^32^ or Trinity^30^. For isoform-sequencing (Iso-Seq) of full-length transcripts, cDNAs were prepared from polyA+RNA using SMARTer® PCR cDNA Synthesis Kit (Takara Bio Inc., Shiga, Japan). The cDNA samples were size-fractionated with the BluePippin™ system (Sage Science, MA, USA) and 700–1500 bp, 1500–3000 bp and 3000–6000 bp fractions were analysed with a PacBio®RSII sequencer (Pacific Biosciences, CA, U.S.A.). Iso-Seq-contigs were constructed using the RS_IsoSeq protocol in SMRT Analysis (v2.3.0) with the parameter of estimated cDNA size. The *de novo* assembled RNA-seq-contigs and Iso-Seq-contigs were mapped to the genome by BLAT^33^, and each contig was annotated when at least 80% of its sequence was mapped. The mapped transcript data were used to make longer transcript models with PASA2 v. 2.0.2^34^.

### Annotation of repeat sequences

Repeat sequences of *P. micropora* were identified using the RepeatModeller package (v. open-1-0-8, http://www.repeatmasker.org) and masked by RepeatMasker v. 4.0^27^, using the identified-model repeat sequences and the repeat sequences of Repbase (ver. 20150807)^35^ (https://www.girinst.org/repbase/). The model repeat sequences were annotated based on the sequence homology search against the NCBI nr database by BLASTX^36^. Representative Polintons and retrotransposons in Supplementary Table 3 were identified by manual inspection of the *P. micropora* draft genome aided by a sequence similarity search using BLAST software^36^.

### Gene annotation

Protein genes of *P. micropora* were annotated by a combination of transcriptome-based gene modelling, *ab initio* gene prediction and protein homology-based gene prediction. In the transcriptome-based gene modelling, the transcripts were masked first by RepeatMasker because many spurious repetitive sequences, which were not removed by genome-repeat-masking, were detected. We discarded the transcript models when more than 80% of the region was masked by RepeatMasker. Exceptions were made when ORFs (> 50 aa) were predicted from the unmasked region. The ORFs and coding sequences of transcripts were predicted with the Transdecoder Utility of Trinity^37^. *Ab initio* gene prediction was performed by Augustus^38^, whose training was performed using Iso-Seq data. Since *P. micropora* protein genes often contain simple repeat sequences in the exon regions, we avoided masking them for the *ab initio* gene prediction. The protein homology-based gene prediction was performed by Exonerate^39^ after masking both the simple repeat and the interspersed repeat sequences, because the homology search in the presence of simple repeat sequences generated an extraordinary number of meaningless candidates (data not shown). In the Exonerate analysis, we used the protein sequences that passed the prescreening by BLASTX search (*P. micropora* genome vs. a local protein database composed of Uniprot and 4 rhizarian organisms, *B. natans*, *P. brassicae, P. chromatophore* and *R. filosa*). All gene models described above were combined, and the best one for each gene locus was chosen according to the bit score of the BLAST search, the presence/absence of transcript and ORF length. The quality of the genome assembly and the annotation was assessed by BUSCO v. 1^40^.

### Detection of DNA virus-like fragments

Genomic segments with discernible boundaries and distinct from the rest of the genome by the following four criteria were denoted as DNA virus-like fragments or putative DNA virus fragments: 1) repeat sequences detected by RepeatModeller but distinct from retrotransposons and the Maverick/Polinton-type transposons, 2) higher GC content, 3) large heterogeneous intron-less gene clusters and 4) ORFs similar to DNA virus proteins are encoded; DNA virus genes are found within the top 100 by BLASTP search against the NCBI nr database. The detailed procedure is as follows. Genomic scaffolds containing repeat sequences of the unknown class^27^ were subjected to ORFfinder^41^, and ORFs detected were annotated by BLASTP^36^ search against the NCBI nr database. The GC% distribution was analysed with CLC genomic workbench 7.0.3 (Qiagen).The virus copy number was analysed using the BLASTN^36^ program searching the simple repeat-masked draft genome for virus coding sequences (Supplementary Table 3) or 100 b fragments generated by slicing the virus-containing scaffolds (Fig. 3, Extended Data Fig. 6). BLASTN-redundant hits were manually removed.

### GO-term-, metabolism-pathway-, orthogroup- and protein-domain analysis

GO-terms were acquired by InterproScan 5^42^, and the enrichment analysis was performed by web-based GOstat^43^ (http://gostat.wehi.edu.au/) using 27,653 GO-terms of 12,007 *P. micropora* genes and 88,634 GO-terms of 39,773 genes of 4 rhizarian organisms (*B. natans, P. brassicae, R. filosa* and *P.micropora,*). Metabolism-pathways were analysed using KAAS of KEGG^44^ (https://www.genome.jp/kegg/kaas/). Orthogroup-analysis of Extended Data Fig. 3 was performed by Orthofinder^45^ using a local protein database (Supplementary Table 10). Protein domain information in Extended Data Fig. 7 was acquired by NCBI conserved domain (CD) search (https://www.ncbi.nlm.nih.gov/Structure/cdd/wrpsb.cgi)^46^.

### nuclear-localized plastid DNA (nupDNA) and mitochondria DNA (numDNA) analysis

Using the *P. micropora* plastid genome (DDBJ Accession No. LC490351) and mitochondria genome sequence (DDBJ Accession No. LC490352) for queries, nupDNA and numDNA were searched for by BLASTN against the *P. micropora* draft genome and the raw sequence reads of Illumina HiSeq2500. In the raw read-based nupDNA and numDNA analysis, chimeric segments of the organelle-like and non-organelle sequences, which represent the junction region of nuclear localized organelle DNA, were surveyed. The detected reads were assembled by CodonCode Aligner (CodonCode Corporation, MA, U.S.A.). The chimera artefacts due to sequencing adaptors, or contaminated bacterial and mitochondria genomes, were identified by BLAST analysis using the NCBI database (nt, nr) and the mitochondria genome sequence (LC490352).

### Phylogenetic analysis and the divergent time analysis

Multiple sequence alignment analyses were performed by MUSCLE^47^ and MAFFT^48^. The phylogenetic trees were constructed using MEGA packages (version 6^49^, 7^50^ and -CC^51^) and IQ-tree^52^. The divergent time analysis was performed using the RelTime method^53^ implemented in MEGA 6. Parameters used for phylogenetic analyses are shown in Supplementary Table 6 and 9, respectively.

### Analysis of EGT/HGT candidates

*P. micropora* nuclear genes derived from cyanobacteria are referred to as EGT, and those derived from the rest of the bacteria are defined as HGT. To screen the EGT/HGT candidates, *P. micropora* MYN1 genes were used as queries for the BLASTP search against the NCBI nr database with e-value ≤ 1e^-10^, and top-hitting genes of the *Paulinella* plastid genes or prokaryotic protein sequences were selected. These genes were subjected to multiple alignment analyses by MUSCLE using the protein sequences of a local protein dataset in Supplementary Table 10 and of the best BLASTP hit sequences. The obtained data sets were subjected to a neighbour joining (NJ) phylogenetic analysis (MEGA6, 7, CC) to choose *P. micropora* nuclear genes that form sister groups with prokaryotes or photosynthetic eukaryotes. After fine taxon re-samplings from the NCBI nr database, the NJ analysis selection was conducted again. The selected genes were finally subjected to maximum likelihood (ML) analysis using MEGA packages (MEGA6, 7, CC). *P. micropora* nuclear genes that satisfied at least one of the following three criteria were used as EGT and HGT candidates. 1) BLAST analysis of the gene gave no hint of eukaryote genes in the NCBI nr database. 2) EGT and HGT are supported by ML phylogenetic analysis with a high bootstrap value. We adopted a bootstrap value of 95 as threshold when phylogenetically available protein alignment sequence positions were long enough (≥ 100 aa). We lowered the threshold to 70 when the available sites were less than 100 aa. 3) The gene is included in the clade of photosynthetic organisms. To confirm the validity of the above mentioned EGT candidate selection, we also screened the EGT candidates by another independent procedure. The EGT candidates obtained by these two independent analyses were almost overlapping and, therefore, used for the analysis (Supplementary Table 6). The alternative analysis procedure is as follows. We performed a phylogenetic analysis using the software of multiple alignment (MAFFT) and ML phylogenetic analysis (IQ-tree), and alignment trimming tools (trimAI^54^, BMGE^55^). We selected genes hitting alpha-type cyanobacteria^56,57^ within the top 100 by BLAST search against a custom database, which consists of the NCBI nr database supplemented with the protein sequences of 14 phylogenetically informative protists (Supplemental Table 10).

### Estimation of EGT/HGT timing

To estimate the timing of EGT/HGT, we used *P. micropora* genes whose counter genes of *P. chromatophora* CCAC0185 were reported as EGT/HGT candidates^58,59^. In addition, we restricted the analysis to *Paulinella* ortholog’s pairs that form a monophyletic sister group with a high bootstrap value (≥ 70). The nearest protein sequences of HGT/EGT candidates were surveyed from NCBI nr database by BLASTP analysis. Within the top 2000 sequences of BLASTP hits, the phylogenetically nearest gene sequences were estimated using NJ and ML phylogenetic analysis. To estimate gene transfer timing, the branching time point when *P. micropora* MYN1 separated from the nearest non-rhizarian organisms in the ML phylogenetic tree was calculated using the RelTime method^55^. We used an estimated value of the divergence of *P. micropora* and *P. chromatophora* (45.7–64.7 MYA) based on the 18S rRNA phylogenetic tree corrected by fossil information^11,60^.

### Analysis of Mavericks/Polinton transposons

*P. micropora’s* Mavericks/Polintons transposons were detected from the draft genome by tBLASTN using the sequence of the DNA polymerase (DNA-pol) domain. Thousands of DNA polymerase ORFs, predicted from the genome sequences by Transdecoder^37^, were subjected to NJ phylogenetic analyses. We grouped highly similar copies. Representative sequences that have long ORFs and less ambiguous amino acid residues were selected from each group and subjected to ML phylogenetic analysis.

### Divergent time analysis of the mobile elements

In the divergent time analysis of Marvericks/Polintons DNA polymerases, representative DNA-pol sequences of *P. micropora* MYN1 Polintons having the traits of recent amplification (nucleotide sequence identity between the copies ≥ 90%) were used. For the DNA-pol sequences of *P. chromatophora* CCAC0185 Polintons, the genome sequence-reads (SRR3217293.sra, SRR3217303.sra) were searched by BLASTX (e-value < 1e^-20^) using the sequence of *P. micropora* MYN1 Polintons, and then, the hit-reads were assembled into contigs with CLC Genomic Workbench 7.0 (Qiagen). For the analysis of virus-type GPCR genes, in addition to *P. micropora* MYN1 genes and *P. micropora* putative virus genes, the translated ORFs of *P. chromatophora* CCAC0185 transcripts were analysed. The GPCR gene family was detected by Orthofinder^45^ and BLASTP^36^ search. In this analysis, the genes encoding seven intact trans-membrane domain sequences, of which all seven trans-membrane helices can be identified by CD search^46^, were used. Furthermore, several genes predicted *ab initio* by Augustus without any supporting experimental data were removed from the analysis, because their gene models appeared to be artificial from applying eukaryotic splicing rules to virus-like fragments. The divergent time points of the mobile elements and GPCR genes were calculated by setting the nearest branching point of *P. micropora* MYN1 and *P. chromatophora* CCAC0185 at 45.7–64.7 MYA.

### Statistical analysis

In GO-term enrichment analysis, Fisher’s exact test (two tailed test) was performed and the p-values corrected with false discovery rate (Benjamini) were calculated. In the phylogenetic analysis, bootstrap test with ≥100 replicates was performed.

## Acknowledgements

We thank Dr. Terabayashi T. for his kind instructions in SEM analysis. Computations were partially performed on the NIG supercomputer at ROIS National Institute of Genetics. This work was supported by JSPS KAKENNHI grants: 221S0002, 15K14554, 1 and 6K14788, and by a Grant-in-Aid for Scientific Research on Innovative Areas 3308 of the Ministry of Education, Culture, Sports, Science and Technology of Japan.

## Author contributions

M. M. prepared the *P. micropora* MYN1 genome samples, identified the putative virus sequence in the *P. micorpora* MYN1 genome and wrote the draft manuscript. M. M. and A. K. performed the RNA sample preparation. Y. M., H. N., A. T. and A. F. performed the genome sequencing, the genome assembly and the Iso-Seq analysis. Y. S. performed the RNA-seq sequencing analysis. M. N. and K. I. cultured *P. micorpora* MYN1. M. M., A. K., M. T., H. N., H. T., S. S., M. N. and K. I. annotated the *P. micropora* MYN1 genome. M. M., T. N. and R. K. performed the phylogenetic analysis. T. N., R. K. and Y. I. analysed the organelle (chromatophore, mitochondria) genome. J. O. organized and managed the *P. micropora* MYN1 genome project and finalized the manuscript. We thank the members of Comparative Genomics Laboratory in National Institute of Genetics for technical and computational assistance.

## Extended Data legends

**Extended Data Fig. 1.**
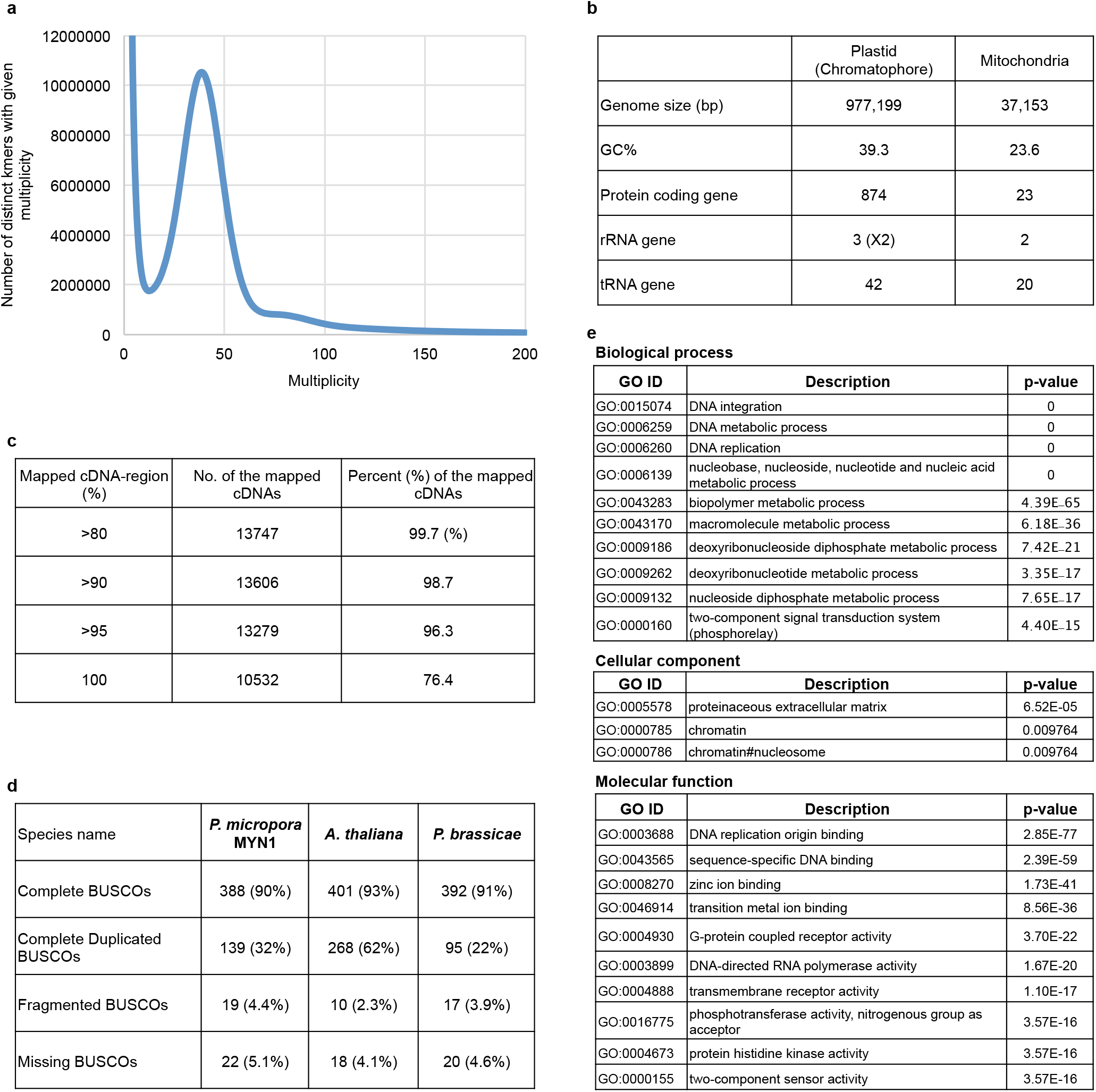
Information of the *P. micropora* MYN1 genome. **a**, Estimation of the *P. micropora* MYN1 genome size by K-mer frequency analysis. Single peak at 39 of the multiplicity are detected with 31-mer, utilizing 52,592,411,100 b from Illumina 500b pair-end reads. From the peak value and the used reads length, 1.35 Gb genome size was estimated. **b**, Summary of the *P. micropora* MYN1 organelle genome. **c**, Validation of the genome assembly by mapping of the sequences of the isoform sequencing (Iso-Seq) transcripts. 13787 non-redundant Iso-Seq sequences of the intron-containing genes, either hitting protein sequences of the Swiss-Prot database by BLASTX search (e-value ≤ 1e^-60^) or containing long ORFs (≥ 300 amino acids), were mapped on the draft genome. **d**, Assessment of the genome assembly using 429 BUSCO v. 1 (Benchmarking Universal Single-Copy Orthologs) genes. In comparison with other eukaryote genome assemblies, the BUSCO values of a photosynthetic organism (*Arabidopsis thaliana*) and that of the well-assembled genome of a rhizarian organism (*Plasmodiophora brassicae*) are shown. **e**, The top 10 GO-terms that are significantly overrepresented in the *P. micropora* MYN1 genome compared with other rhizarian organisms (*Bigelowiella natans*, *Plasmodiophora brassicae*, *Reticulomyxa filosa*). Fisher’s exact test p-values corrected with false discovery rate (Benjamini) are represented.

**Extended Data Fig. 2.**
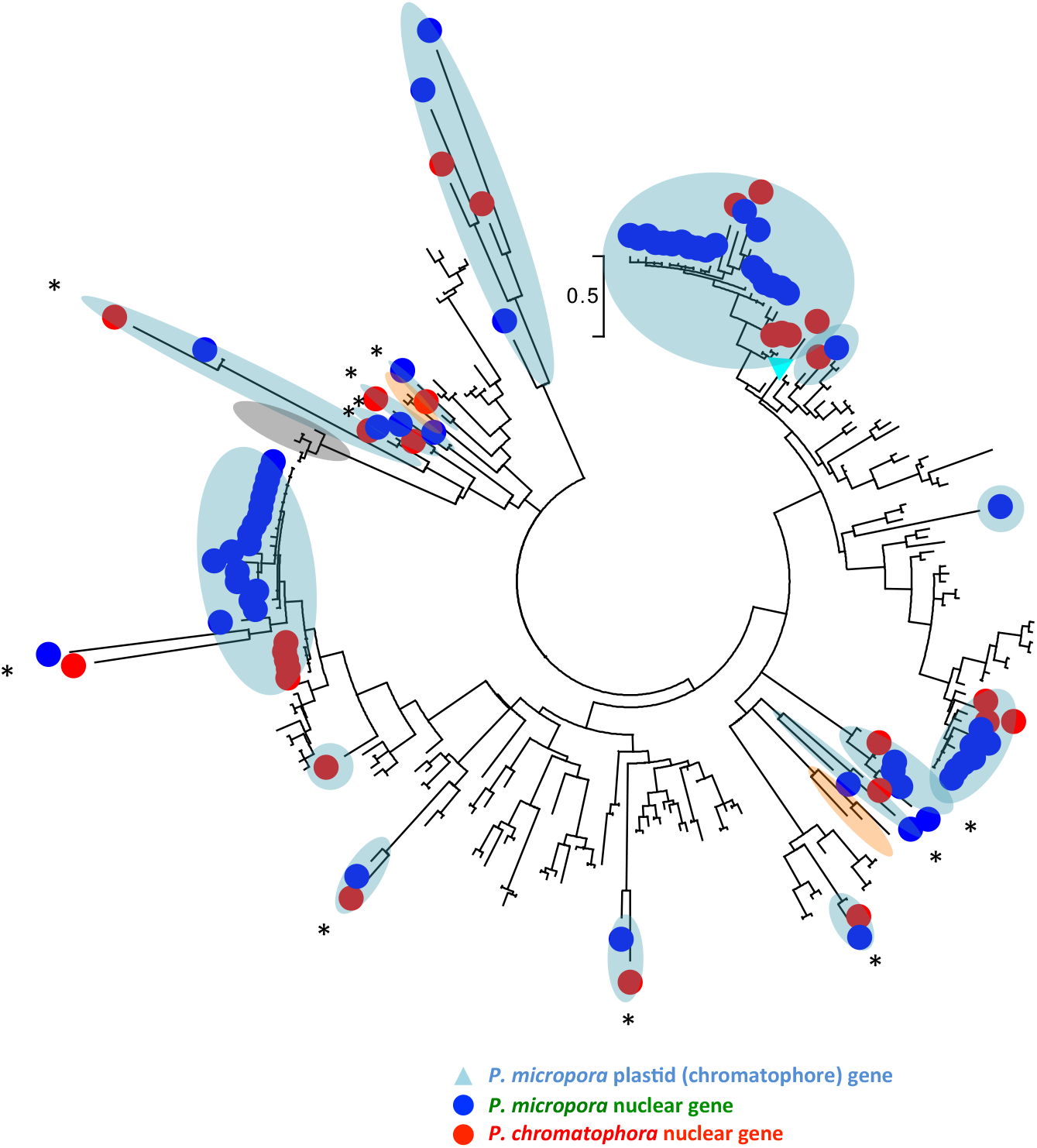
A ML phylogenetic tree of Hlips of *P. micropora* MYN1 and *P. chromatophora*. Hlips of cyanobacteria, photosynthetic eukaryotes and cyanophages are used as a reference for operational taxonomy units. Sixty-four *P. micropora* Hlip genes were subjected to phylogenetic analysis and grouped according to the clade. The redundant paralogs of identical sequences were removed. Asterisks: *Paulinella* ortholog-pair supported by a bootstrap value >70. Blue and red circle: nuclear encoded Hlip of *P. micropora* and *P. chromatophora*. Light-blue triangle: *P. micropora* plastid (chromatophore) Hlip. Phylogenetic branch of the *Paulinella* gene (light blue), virus (grey) and eukaryote Hlip-like gene (orange) are highlighted.

**Extended Data Fig. 3.**
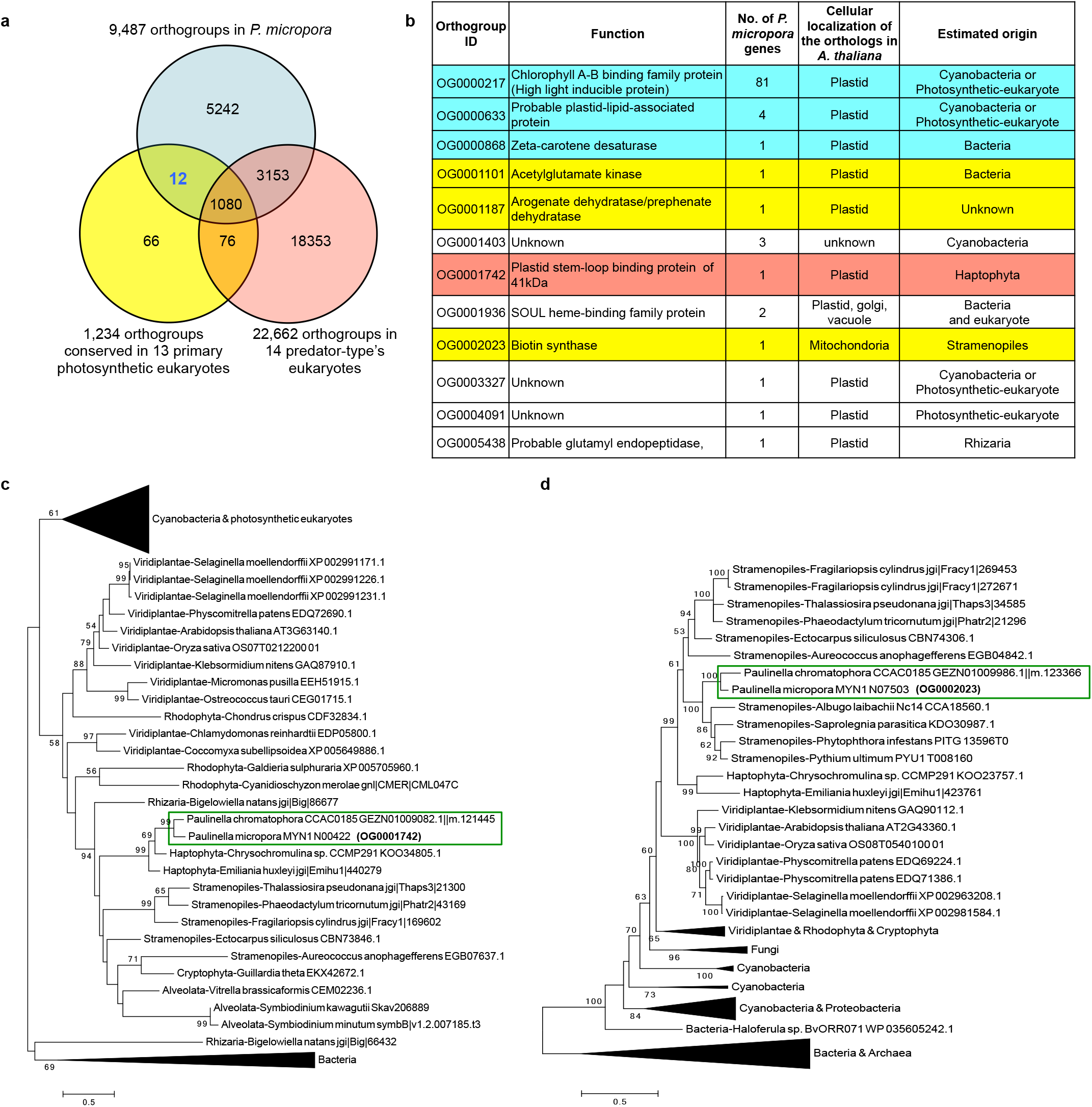
Comparison of the orthogroups of *P. micropora*, primary photosynthetic eukaryotes and predator-type eukaryotes. **a**, Venn diagram of *P. micriopora* orthogroups with the predator-type eukaryote orthogroups and the conserved orthogroups in primary photosynthetic eukaryotes. Orthogroups were detected by Orthofinder from the gene sets of *P. micropora* MYN1, 14 predator eukaryotes (*Acanthamoeba castellanii*, *Dictyostelium discoideum*, *Entamoeba histolytica HM-1*, *Bodo saltans*, *Naegleria gruberi*, *Caenorhabditis_elegans*, *Drosophila melanogaster*, *Monosiga brevicollis MX1*, *Oxytricha trifallax, Paramecium tetraurelia*, *Stylonychia lemnae*, *Tetrahymena thermophila*, *Reticulomyxa filosa*, *Thecamonas trahens ATCC50062*) and 13 photosynthetic eukaryotes (*Arabidopsis thaliana*, *Chlamydomonas reinhardtii*, *Coccomyxa subellipsoidea C-169*, *Klebsormidium nitens*, *Micromonas pusilla CCMP1545*, *Oryza sativa*, *Ostreococcus tauri*, *Physcomitrella patens*, *Selaginella moellendorffii*, *Chondrus crispus*, *Cyanidioschyzon merolae*, *Galdieria sulphuraria*, *Cyanophora paradoxa*). **b**, Annotations of 12 orthogroups conserved in primary photosynthetic eukaryotes, but not in predator eukaryotes. The functional annotation and the cellular localization were based on UniprotKB information of *Arabidopsis thaliana* orthologs, and manual annotations are represented in parenthesis. The estimated origins were based on the ML phylogenetic analysis. Genes involved in light acclimation (cyan), nutrient auxotrophy (yellow) and organelle gene expression (magenta) are highlighted. **c** and **d**, ML phylogenetic trees of orthogroup genes acquired by HGT from other eukaryotes; Haptophyta (**c**, OG0001742) and Stramenopiles (**d**, OG0002023). The phylogenetic analysis of **c** and **d** were performed using a LG+G (**c**) and a LG+G+I model (**d**), respectively. Photosynthetic *Paulinella* species are boxed.

**Extended Data Fig. 4.**
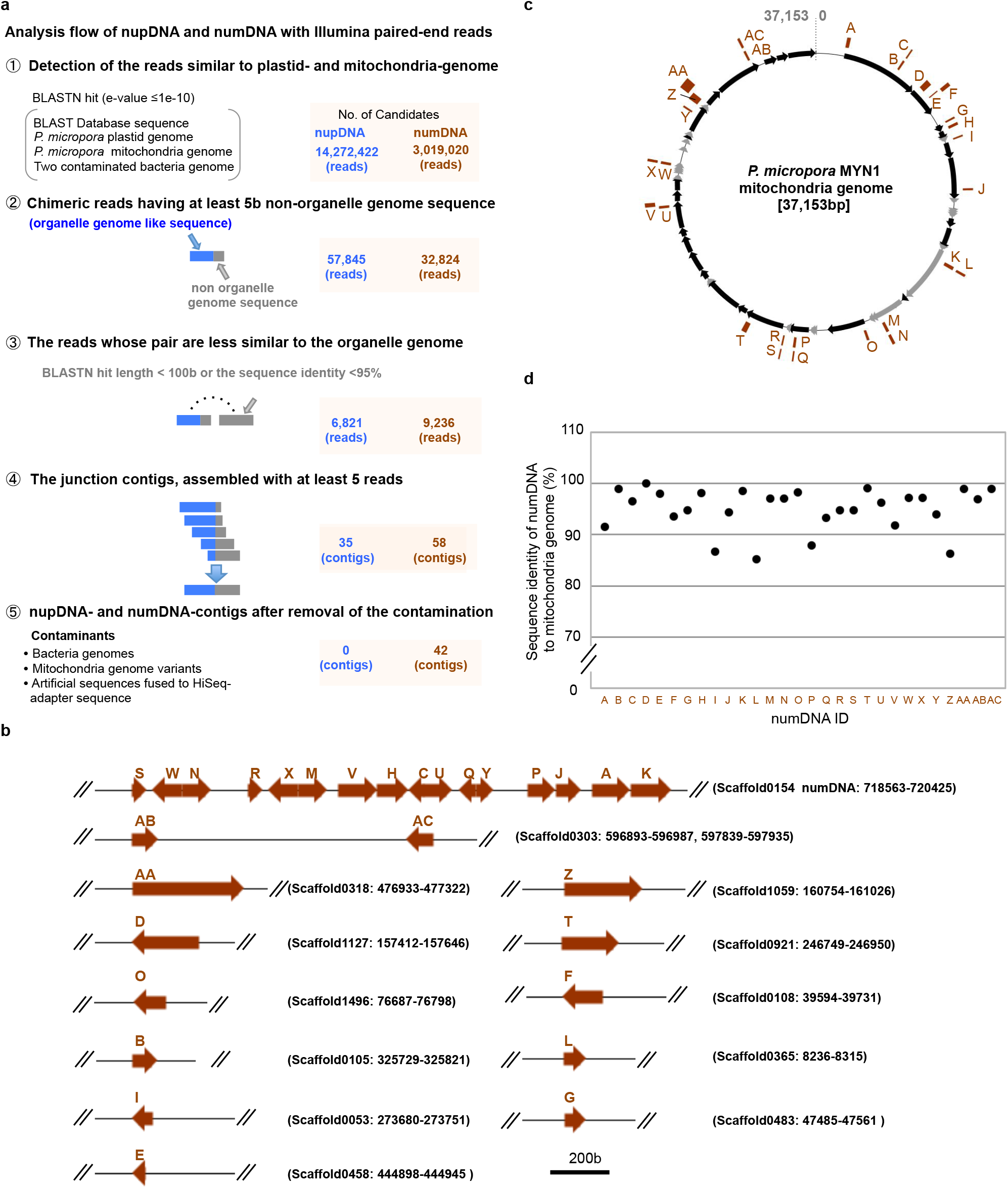
Analysis of nuclear-localized plastid DNA (nupDNA) and mitochondria DNA (numDNA) in *P. micropora*. **a**, Analysis of the junction sequences of nupDNA and numDNA using Illumina raw-reads sequences. 902,121,079 quality-trimmed HiSeq paired-end sequence reads (insert size: 300 b, 500 b) were analysed. **b**, numDNA sequence structures in the *P. micropora* draft genome. **c**, Distribution of numDNA sequences in mitochondrial genome positions. **d**, Nucleotide percent identity of numDNA compared with the mitochondria genome sequence. NumDNA directions are represented by right and left arrows which denote clockwise- and anti-clockwise directions of the mitochondria genome sequence (**c**), respectively.

**Extended Data Fig. 5.**
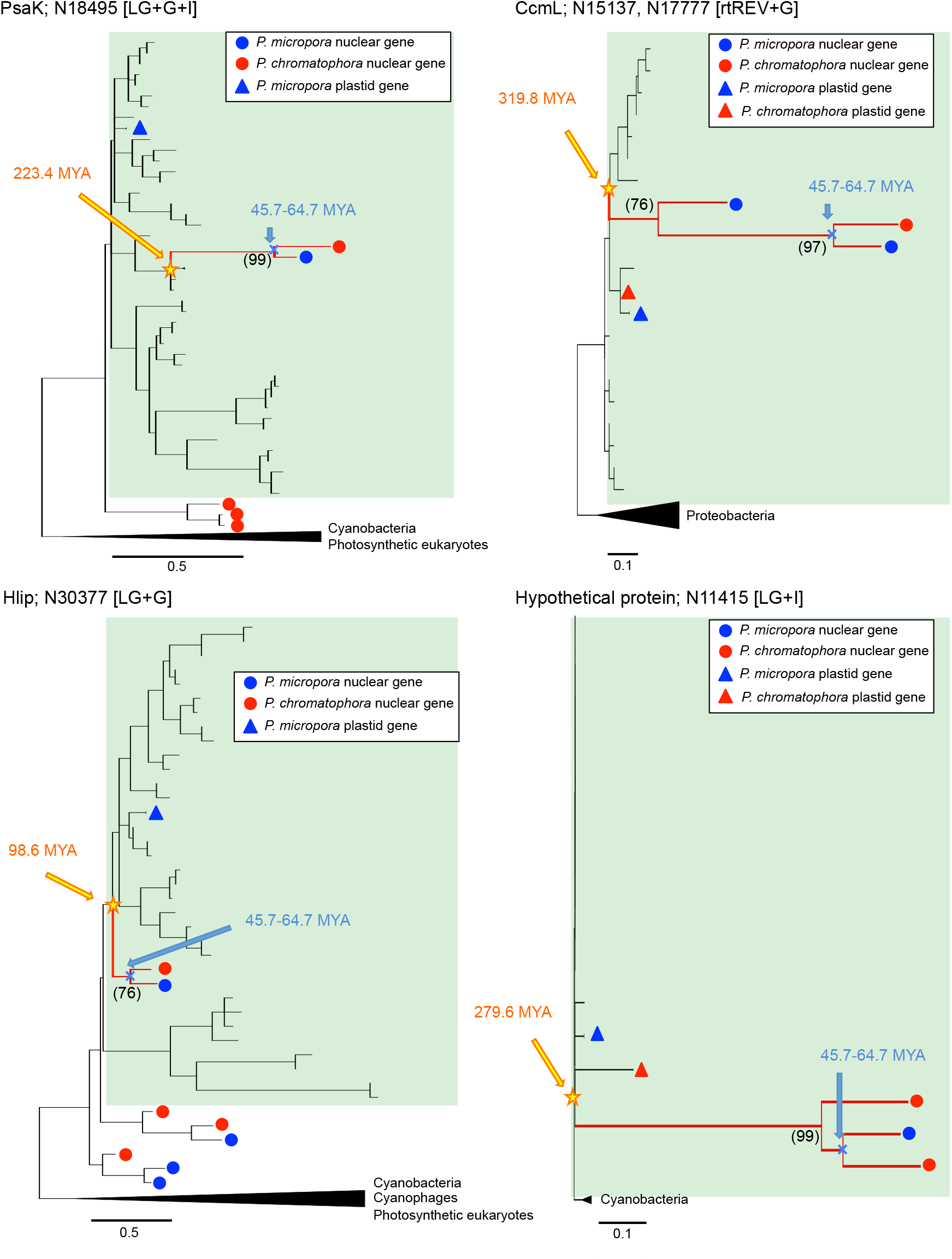
ML phylogenetic trees of four EGT candidates whose counterpart orthologs are found in the plastid (chromatophore) genome of *P. micropora*. The clades of the *Paulinella* plastid and cyanobacteria are highlighted in green. Blue circle: *P*. *micropora* nuclear gene, red circle: *P. chromatophora* nuclear gene, blue triangle: *P. micropora* plastid gene, red triangle: *P. chromatophora* plastid gene, unmarked: cyanobacterium gene. The branch of *P. micropora* and *P. chromatophora* nuclear genes supported by high bootstraps (numbers in the parenthesis) is represented as a red line. The divergence time of *Paulinella* nuclear genes from the plastid- or other cyanobacterial-genes (yellow stars) were estimated by RelTime methods of MEGA6 using the divergent time of *P. micropora* and *P. chromatophora* (45.7–64.7 MYA) (blue cross). Psak: Photosystem I subunit K, CcmL: CO_2_ concentrating mechanism protein, Hlip: High light inducible protein. Brackets mean the substitution model used in the phylogenetic analysis. Trees with species names are available at the repository (https://figshare.com/s/a665678c48d0af073894).

**Extended Data Fig. 6.**
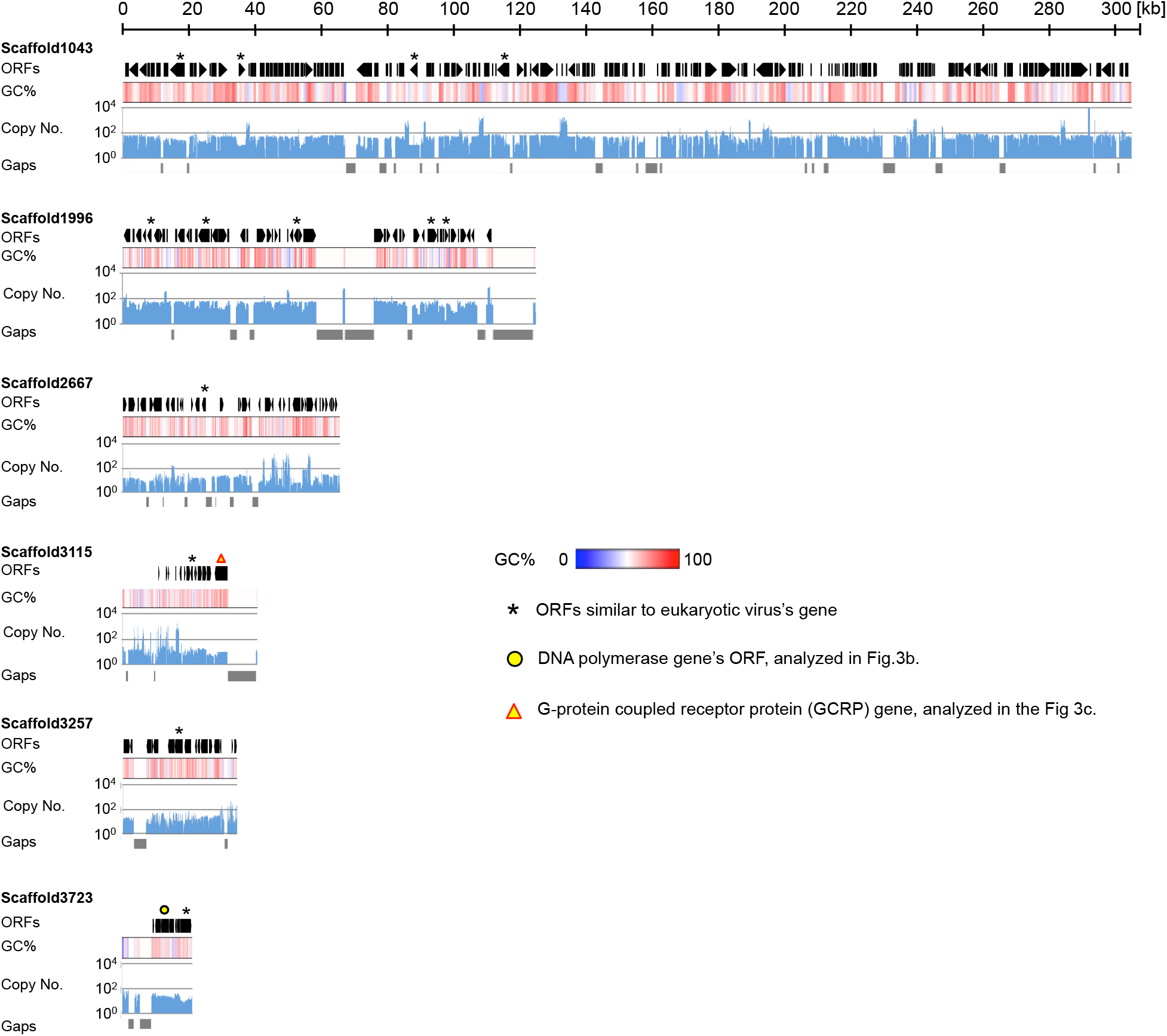
Putative DNA virus fragments detected in the *P. micropora* draft genome. ORF structures, GC%, copy number and sequence gaps are represented as described in the legend to Fig. 3.

**Extended Data Fig. 7.**
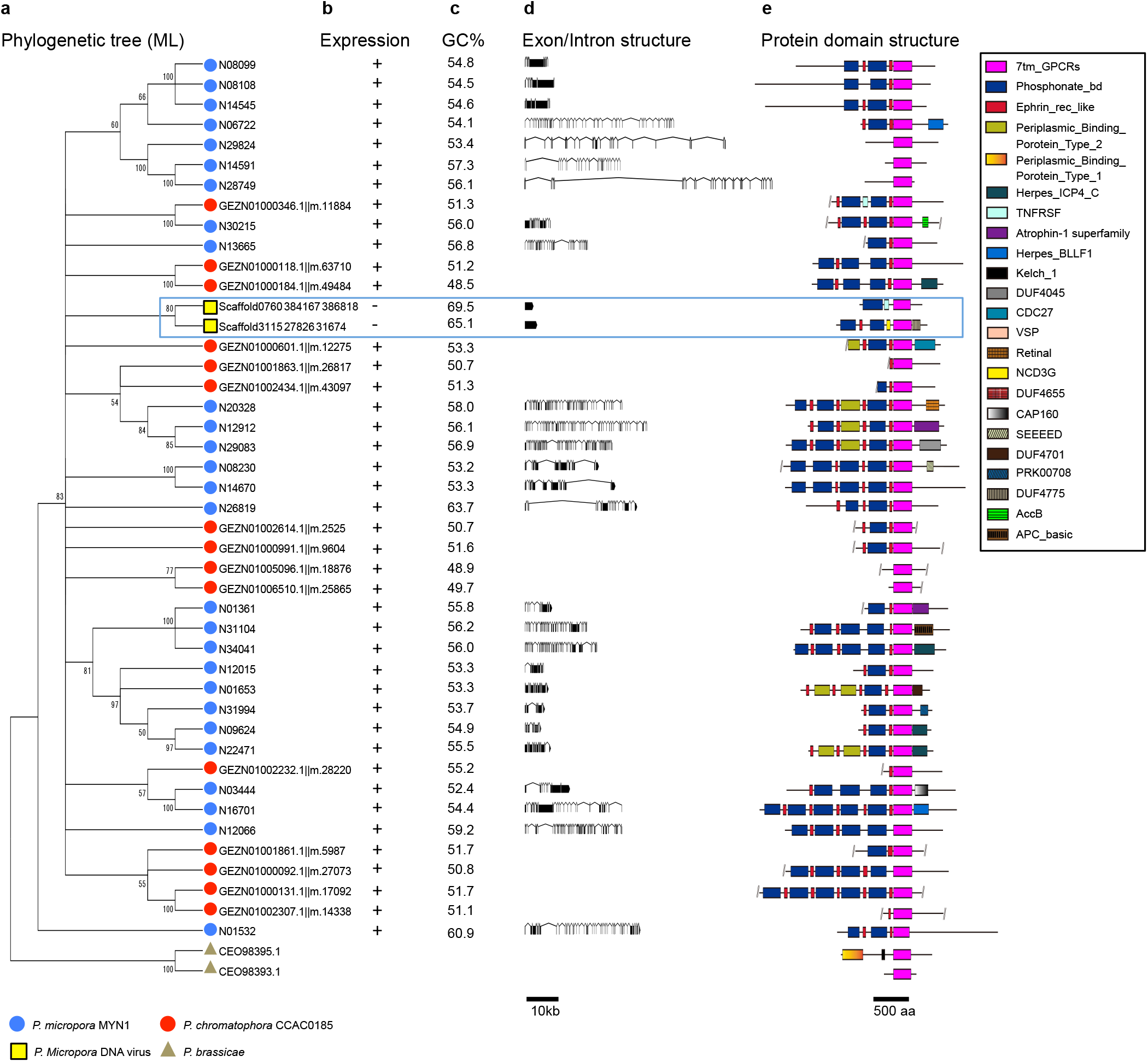
Characteristics of virus-type GPCRs of *P. micropora* and *P. chromatophora.* **a**, A ML phylogenetic tree of seven transmembrane domains of GPCRs. GPCRs in *P. micropora, P. chromatophora, P. brassicae* (CEO98393.1, CEO98395.1), and those detected in DNA viral fragments were analysed using a LG+G+F model. Branches with bootstrap values <50 are condensed. **b**, Existence of the transcript. **c**, GC% of the coding sequence of GPCR genes. **d**, Exon/intron structures of the genes. **e**, Protein domain structures. Protein domains were detected by CD search^48^. GPCRs in putative DNA viruses are boxed with a blue line.

**Extended Data Fig. 8.**
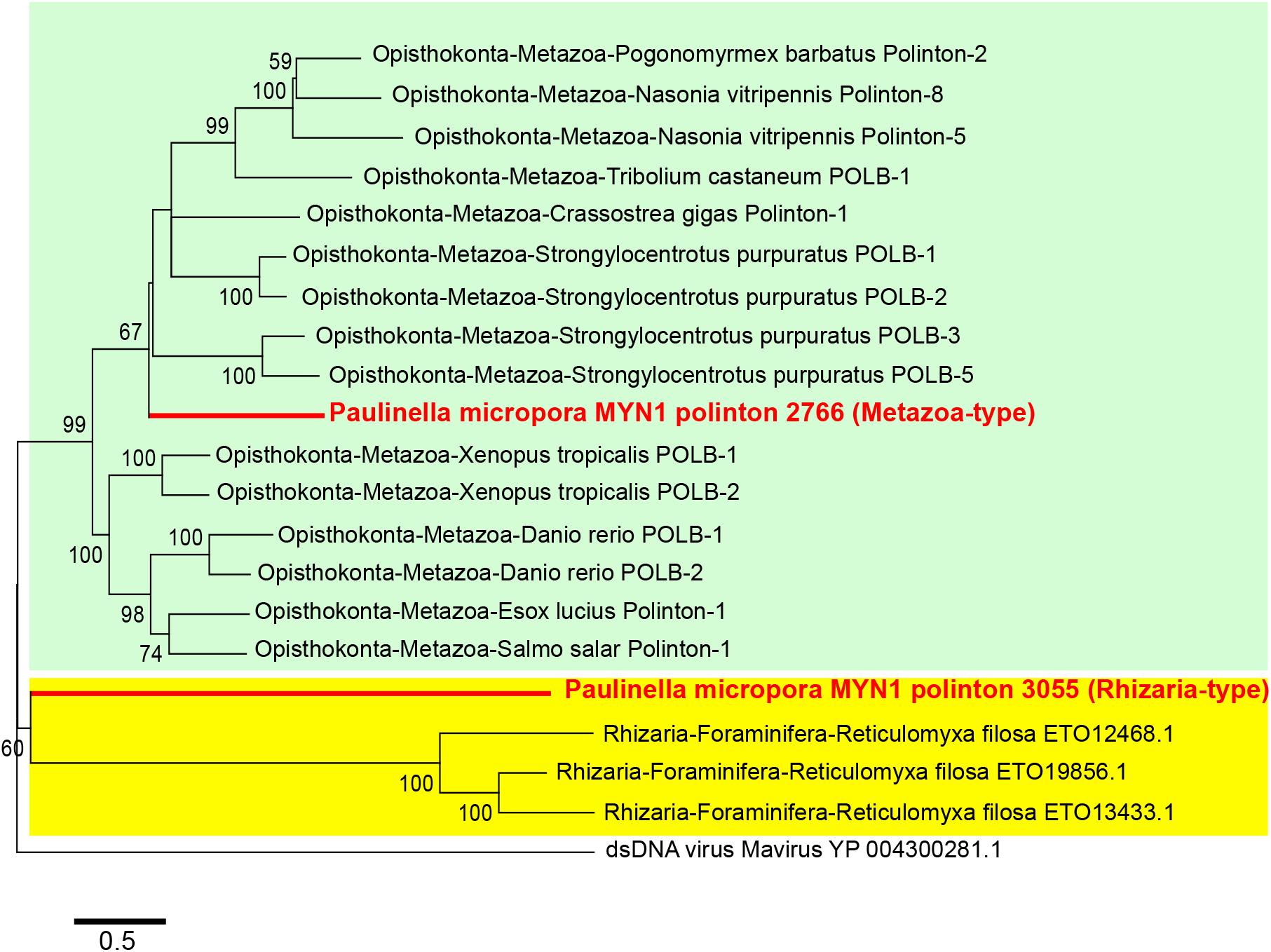
ML phylogenetic tree of DNA polymerases of Polintons. **a**, DNA polymerase sequences of *P. micropora* Polintons (No. 2766 and No. 3055), *R. filosa’* Polintons (Genbank, ETO12468.1, ETO19856.1, ETO13433.1), metazoan Polintons in Repbase^37^ and Mavirus (Genbank, YP 004300281.1) were analysed with a LG+G+I model. The metazoan and rhizarian groups are highlighted in green and yellow, respectively.

## Supplementary information

**Supplementary Table 1. Summary of the raw sequence data of the *P. micropora* genome.**

**Supplementary Table 2. Annotation and categorization of the *P. micropora* RepeatModeller sequences.** *The categorization is based on the sequence similarities with the manually curated repeat sequences in Supplementary Table 3 by BLASTX search (e-value < 1e^-5^).

**Supplementary Table 3. Annotation of the repeat elements manually identified from the *P. micropora* draft genome.** *The copy number of the repeat elements was estimated by BLASTN search against the *P. micropora* genome. Redundant BLASTN hits at the same genome locus were manually removed.

**Supplementary Table 4. *P. micropora* MYN1 gene list.**

**Supplementary Table 5. GO-terms over- and under-represented in the *P. micropora* MYN1 genome compared with the genome of other rhizarian organisms (*B. natans, P. brassicae and R.filosa*).** Significantly enriched GO-terms with Fisher’s exact test p-value less than 0.01 are represented.

**Supplementary Table 6. EGT and HGT candidates in *P. micropora* MYN1.** Genes satisfying at least one of the following criteria were considered as EGT and HGT candidates. 1) No hits to eukaryote genes of the NCBI nr database by BLAST analysis* 2) EGT and HGT are supported by ML phylogenetic analysis with a high bootstrap value (≥95) except when the phylogenetically available protein alignment sequences were short (< 100 amino acids)** 3) *P. micropora* genes were embedded in the clade of photosynthetic organisms in the phylogenetic tree. * BLASTP top 1000 hits (NCBI nr, e-value < 1e^-10^) were classified. A: Archaea, B: Bacteria, E: Eukaryotes, V: Viruses, O: Others, U: Unknown. ** We lowered the threshold of the bootstrap value to 70 when the alignment sequence positions available for the phylogenetic analysis were less than 100 amino acids.

**Supplementary Table 7. Detection of P. *micropora* Polintons by tBLASTn search using DNA polymerase domain sequences.** A tBLASTn search was performed against the *P. micropora* draft genome (e-value < 1e^-10^). The draft genome sequences and the query sequences are available at the repository (https://figshare.com/s/a665678c48d0af073894).

**Supplementary Table 8.** Accession numbers of P. micropora MYN1 genome sequences.

**Supplementary Table 9. Parameters of the ML phylogenetic analysis by MEGA6.**

**Supplementary Table 10. Sequence data used in this study.**

